# Population sequencing for diversity and transmission analyses

**DOI:** 10.1101/2024.06.18.599478

**Authors:** Talima Pearson, Tara Furstenau, Colin Wood, Vanessa Rigas, Jason Sahl, Sara Maltinsky, Bart J Currie, Mark Mayo, Carina Hall, Paul Keim, Viacheslav Fofanov

**Author notes:** Corresponding Author: Talima Pearson.

## Abstract

Genomic diversity in a pathogen population is the foundation for evolution and adaptations in virulence, drug resistance, pathogenesis, and immune evasion. Characterizing, analyzing, and understanding population-level diversity is also essential for epidemiological and forensic tracking of sources and revealing detailed pathways of transmission and spread. For bacteria, culturing, isolating, and sequencing the large number of individual colonies required to adequately sample diversity can be prohibitively time-consuming and expensive. While sequencing directly from a mixed population will show variants among reads, they cannot be linked to reveal allele combinations associated with particular traits or phylogenetic inheritance patterns. Here, we describe the theory and method of how population sequencing directly from a mixed sample can be used in conjunction with sequencing a very small number of colonies to describe the phylogenetic diversity of a population without haplotype reconstruction. To demonstrate the utility of population sequencing in capturing phylogenetic diversity, we compared isogenic clones to population sequences of *Burkholderia pseudomallei* from the sputum of a single patient. We also analyzed population sequences of *Staphylococcus aureus* derived from different people and different body sites. Sequencing results confirm our ability to capture and characterize phylogenetic diversity in our samples. Our analyses of *B. pseudomallei* populations led to the surprising discovery that the pathogen population is highly structured in sputum, suggesting that for some pathogens, sputum sampling may preserve structuring in the lungs and thus present a non-invasive alternative to understanding colonization, movement, and pathogen/host interactions. Our analyses of *S. aureus* samples show how comparing phylogenetic diversity across populations can reveal directionality of transmission between hosts and across body sites, demonstrating the power and utility for characterizing the spread of disease and identification of reservoirs at the finest levels. We anticipate that population sequencing and analysis can be broadly applied to accelerate research in a broad range of fields reliant on a foundational understanding of population diversity.

**Author Summary:** The ability to characterize diversity in a single bacterial population (i.e., a single host or even a single body site) is critical for understanding adaptation and evolution, with far-reaching implications on disease treatment and prevention that include revealing patterns of spread and persistence. While the scientific community has made great strides in sequencing methods to characterize single colonies and entire communities, there is a dearth of studies at the population level. This is because 1) the need to culture and sequence a sufficiently representative number of isogenic colonies is prohibitive, and 2) the theoretical foundation for characterizing a population by sequencing a single sample (as is done for microbiome and metagenomic analyses) has not been developed. Here, we introduce this theoretical foundation and validate its applicability by characterizing a lung infection caused by *Burkholderia pseudomallei*. We also demonstrate the utility of this method in determining the directionality of spread of *Staphylococcus aureus* between people and across body sites within the same host (a level of spatial resolution that has not been previously performed). We anticipate that this work will open the door to a host of new studies and discoveries across a diverse set of microbiological fields.

## Introduction

Genomic analyses of pathogens provide insights into their diversity, history, spread, and transmission, all of which are critical for mitigation. The potential for whole genome sequencing as a tool for determining high-resolution and high-accuracy evolutionary history of single bacterial species was established two decades ago [1]. Initially, high sequencing costs prioritized capturing the breadth of diversity within a species, providing a pathway for addressing broad evolutionary questions by characterizing the genetic repertoire (e.g., core, accessory, and pan-genome) and global phylogeographic pattens. More recently, it became clear that a single colony did not represent the diversity in a population, and now that genome sequencing is routine, fine-scale evolutionary questions reliant on the identification of small genomic differences can be answered. Sequencing multiple representatives of a population to characterize the genetic diversity is critical for answering questions about within-host adaptation and changes of pathogens [2–5], source attribution for microbial forensic investigations [6], and to characterize outbreaks [7–9].

Typically, a small number of genomes from each source is used to characterize diversity to determine infection duration, evolution, and involvement in a transmission chain. However, if there is genetic diversity within the pathogen’s source, inferences based on limited sampling of a population can be erroneous. In such instances, if the organism sampled from a given source is phylogenetically distant from the organism(s) that were transmitted, the source will be erroneously excluded from the transmission chain. Similarly, phylogenetic branching order may lead to false conclusions about persistence and spread as unsampled genomes may have diverged from earlier or later parts of the phylogeny relative to sampled genomes from the same and different sources. Likewise, reliance on temporal data to infer chronological order of infection events may produce misleading results as a recently sampled genome may in fact have a deeper evolutionary origin (a more ancient common ancestor) compared to other genomes that may have been sampled previously. Capturing genomic and phylogenetic diversity at the level of a single host is critical for accurate insights into fine-scale pathogen transmission and spread yet remains a significant challenge.

Characterizing genetic diversity of culturable bacteria may seem straightforward as multiple colonies from a population can be isolated and sequenced. However, you would need to sample almost 60 colonies to have a 95% confidence of capturing a variant present at a 5% frequency (assuming random distribution of minor variant colonies in culture media). Many clinically relevant minority variants are initially present at frequencies much lower than 5%. Sequencing multiple colonies from single sources has provided valuable insights into pathogen population genetics at small-evolutionary scales, but the time and expense required to sequence many samples has led to either a focus on deeply sampling a select case or transmission cluster, or a more cursory examination across multiple cases [4,10–14]. Second-generation sequencing approaches can deeply sample genomic variants in a population, but assessment of diversity is primarily limited to a simple tally of mutations across reads or small windows of the genome [5,15]. For small genomes, linking mutations across reads to reconstruct estimated haplotypes is possible if the population is composed of highly heterogeneous strains with diversity distributed throughout the genome to guide the joining of reads [16]. Third and fourth-generation sequencing methods circumvent the need to stitch reads together for organisms with small genomes (e.g., most RNA viruses). However, even with long reads, the sparsity of variants in larger genomes from relatively homogeneous populations, restrict the joining of reads for haplotype reconstruction.

Here, we show how deep sequencing of a population can be used to characterize the phylogenetic diversity of a pathogen population without haplotype reconstruction or linking mutations across reads. While the full extent of population diversity and phylogenetic branching patterns of variants cannot be characterized with this method, it provides a measure of diversity and defines the phylogenetic boundaries of the population. Diversity and phylogeny are important for understanding the evolutionary history of a single population. For example, an older population will be more diverse than a newly established population (all else constant), but selection and drift will also shape diversity and phylogenetic patterns. Diversity and phylogeny are also critical for understanding interactions between populations. For example, the directionality of spread and transmission can be determined as the recipient population will be less diverse and phylogenetically nested within the diversity of the source. In this work, we: 1) discuss the theoretical foundation for characterizing phylogenetic diversity of a clonal population through population sequencing, 2) explore the utility of population sequencing in capturing the phylogeny of a well-characterized population of *Burkholderia pseudomallei* [4], 3) examine advantages and limitations of population sequencing using a clonal model of population generation and transmission, and 4) use population sequencing to characterize transmission of *Staphylococcus aureus* across persons and body sites within a single person.

## Results

### Genotyping a population of *Burkholderia pseudomallei*

We previously established the phylogeny of 118 clones of *B. pseudomallei* collected over the course of a chronic lung infection between the years 2000-2017 [4]. To this phylogeny, we added 20 sequenced isogenic clones (colonies) from two sputum samples collected on a single day in August 2022 and 2 population sequences from one sputum (sequenced at an average depth of 502 and 931 reads for populations 1 and 2, respectively) by determining the alleles at 162 previously documented SNPs and 22 novel SNPs discovered among the new clones (Figure 1). Sequencing the 20 colonies from this population resulted in 9 novel genotypes located in 3 clades. Surprisingly, the phylogenetic range of the population sequences: 1) did not encompass all of the clone sequences, 2) did not extend beyond the clone sequences, and 3) showed very little overlap (genotypes in the middle clade were almost exclusive to Pop 2 making up approximately 40% of the population and genotypes in the bottom clade were exclusive to Pop 1 making up approximately 30% of the population). For some SNPs, the derived allele was found in very low frequencies, suggesting that for a very small proportion of each population, the phylogenetic range is slightly greater than what is depicted in Figure 1. The 1,441 novel SNPs (458 common between both populations, 269 unique to Pop 1, and 717 unique to Pop 2) found only among the population sequences provide evidence of additional diversity that, without haplotype reconstruction, cannot be placed on the phylogenetic tree. The structuring (uneven distribution) of the population sequences was consistent with the phylogenetic placement of the isogenic clones as the top clade was almost completely composed of clones from Sample 1 and the middle clade was composed of only clones from Sample 2. Although the population sequences were from the same sputum sample, the sample was not homogenized and the portions that made up the two population samples were loops collected at different locations from the viscous sputum matrix. The phylogenetic patterns from 20 isogenic clones and two population sequences provide strong evidence that while both sputum samples contain diverse genotypes, each sputum sample is not well mixed and thus the genotypes are highly clumped suggesting spatial structuring in the lungs.

**Figure 1:**
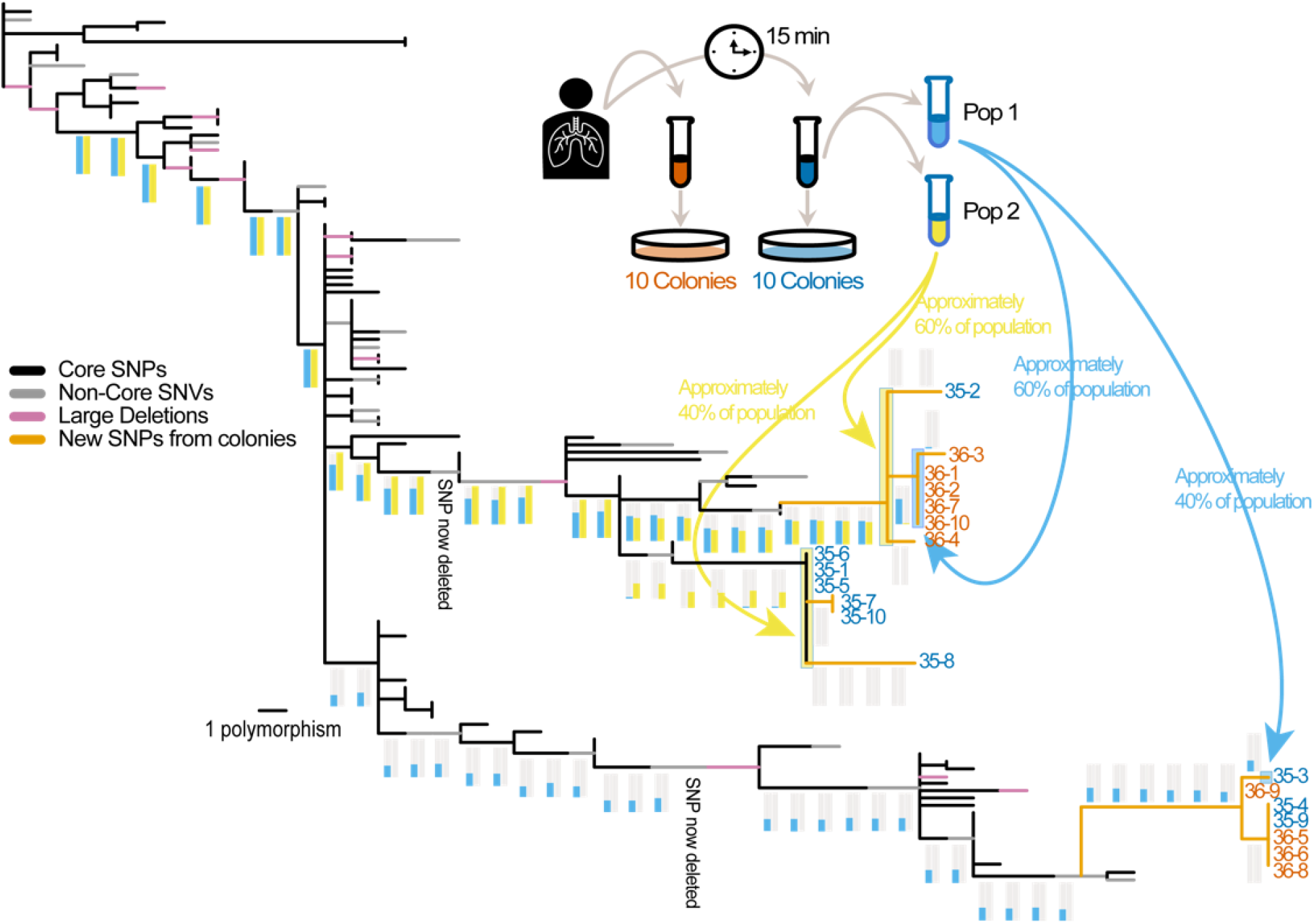
Placement of clonal and population sequences on an established phylogeny of a chronic lung infection of *B. pseudomallei*. Clonal phylogeny showing the previously established phylogeny (black, gray, and pink branches) with new branches (orange) and the location of clones from 2 new sputum samples and population sequences. Experimental overview (inset). Frequencies of the derived SNP alleles from the population sequences along branches leading to the extant populations and clones are shown as histograms with colors indicating the originating population. The estimated phylogenetic range of the content in each population sequence is shown as a colored background to the phylogenetic branches and indicated by blue and yellow arrows. For each population, some SNPs show a very small proportion of the derived allele (e.g., for Pop1 in 3 of the 6 SNPs leading to the middle extant clade) suggesting that while most of the population is contained in the depicted phylogenetic range, a very small portion is not.

### Modeling to determine directionality of transmission/spread

When a wide transmission bottleneck allows for the ecological establishment of a larger number of cells, determination of transmission directionality becomes more challenging, with a decreased likelihood of determining the correct direction, and an increased likelihood of both incorrect or ambiguous results. Of the three methods that we evaluated for determining directionality, Method 3 (phylogenetic entropy) performed better than Methods 1 and 2 (both based on phylogenetic range) (Figure 2). Compared to Methods 1 and 2, although the proportion of correct determinations for Method 3 were fewer with a wider bottleneck, the proportion of incorrect determination were also fewer (Figure 2), resulting in a better net performance. Method 3 was most likely to lead to ambiguous transmission directionality results, but less likely to result in erroneous determinations. The simulations estimated the probability of correctly detecting transmission based on comparing only a single genome from each population and therefore increasing the number of genomes sampled would improve the chances of correctly determining transmission direction. We used Method 3 for further evaluations of the impact of other variables on determination of transmission direction.

**Figure 2:**
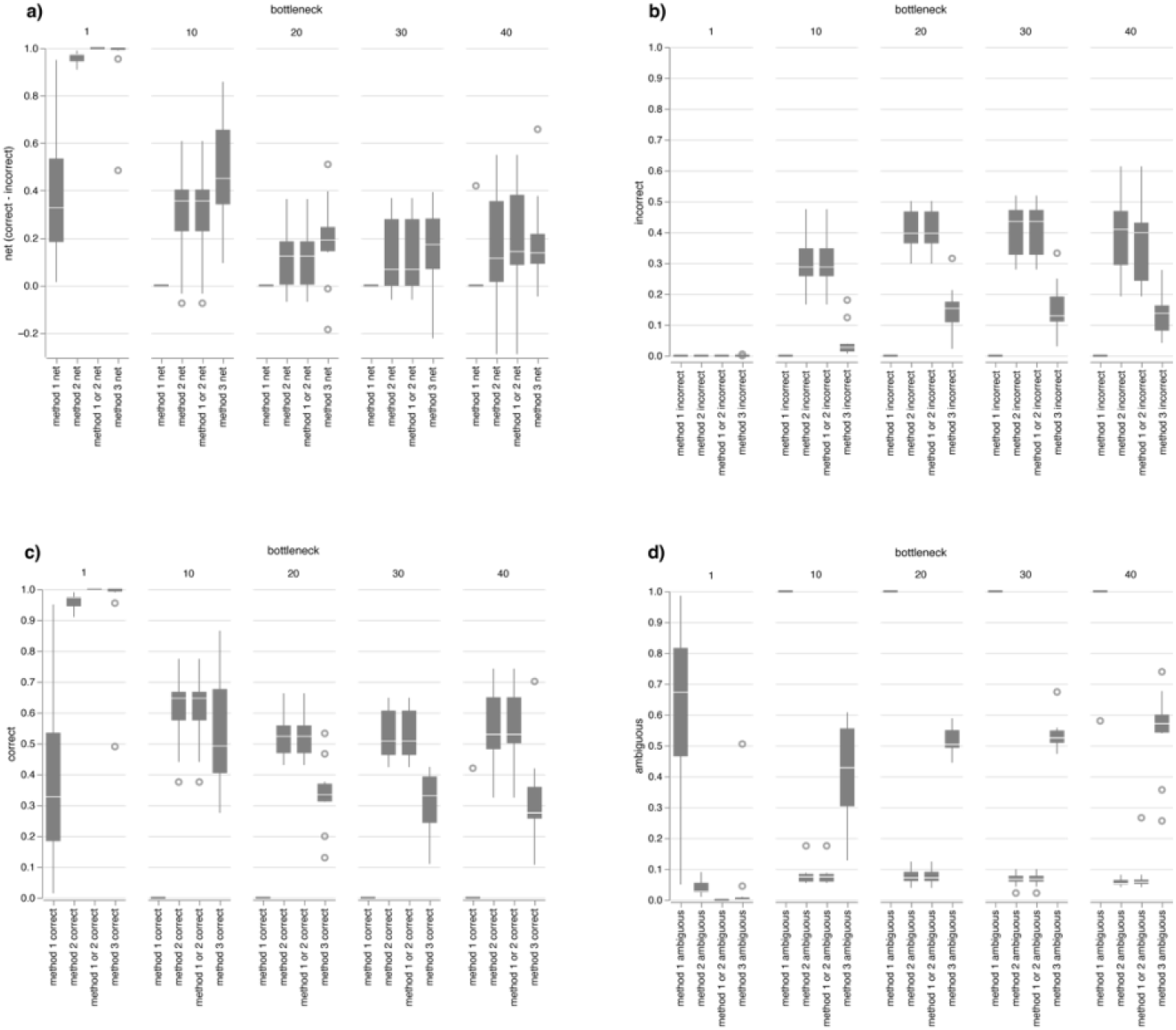
Comparison of three methods to determine transmission directionality. Net proportion of correct results (a) calculated by subtracting incorrect determinations (b) from correct determinations (c). Proportion of ambiguous results (d).

We explored 3 additional variables that are most likely to impact the potential for sequenced populations to yield information on the directionality of transmission. Firstly, as the phylogenetic nesting of the recipient clade(s) within the source clade is necessary for determining transmission directionality, the existence of phylogenetic diversity in the source population before transmission is critical. As evolutionary time, modeled by the number of generations in the source, increases, the likelihood of correct determinations of transmission directionality increases while incorrect and ambiguous determinations decrease (Figure 3a). Secondly, evolutionary time after transmission allows for the source and recipient populations to diverge, reducing the likelihood of correctly determining transmission directionality and increasing the likelihood of ambiguous results (Figure 3b). Thirdly, a wider bottleneck involves the transfer of more cells and potentially more diversity, reducing the likelihood of correct determinations of directionality (Figure 3c). Lastly, selecting more clones for sequencing will increase the likelihood of discovering more branches in the resulting phylogeny and finding additional segregating SNPs that may increase phylogenetic entropy. As such, sampling and sequencing more clones increases the likelihood of correctly determining transmission directionality and decreases the likelihood of ambiguous results (Figure 3d), although this salutary effect diminishes rapidly. It is important to note that for the tested variables, the likelihood of incorrect determination of transmission is small and thus Method 3 provides a conservative estimation of directionality.

**Figure 3:**
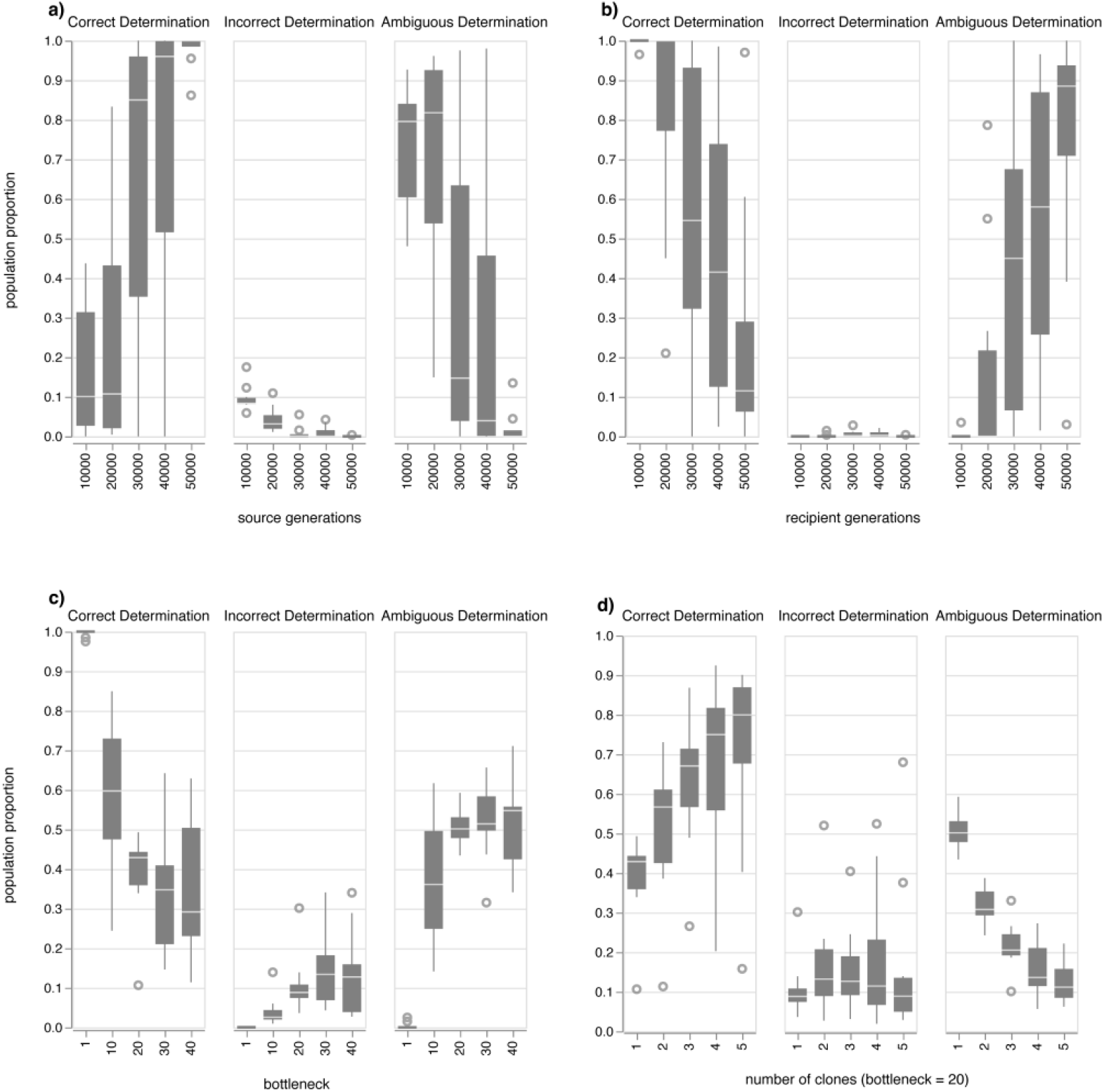
Evaluation of the impact of source generations (a), recipient generations (b), the bottleneck size (c), and number of clones selected for sequencing (d) on determining directionality of evolution. Unless otherwise indicated, simulation parameters were as follows: 100,000 source generations, 10,000 recipient generations, bottleneck size 1, number of clones 1.

### Transmission/spread of *S. aureus*

We next used Method 3 to determine the directionality of transmission and spread of *S. aureus* using empirical examples. Selection of a close outgroup clone for rooting and comparison to clones from each population for SNP discovery is critical for ensuring that SNPs along Branch C are limited to recent mutation events. A more distant outgroup results in more SNPs along Branch C and as more SNP loci are interrogated across the sequenced populations, it becomes increasingly likely that some of those loci will, by chance alone, have segregating alleles. Analyses of SNPs along Branch C must therefore be limited to very recent SNPs and as such, segregating alleles will be the result of shared inheritance rather than spurious independent mutations. Analysis of *S. aureus* genomes from multiple body sites from 4 individuals reveals the potential for population sequencing to identify transmission between persons as well as spread from one site to another within a single person (Figure 4).

**Figure 4.**
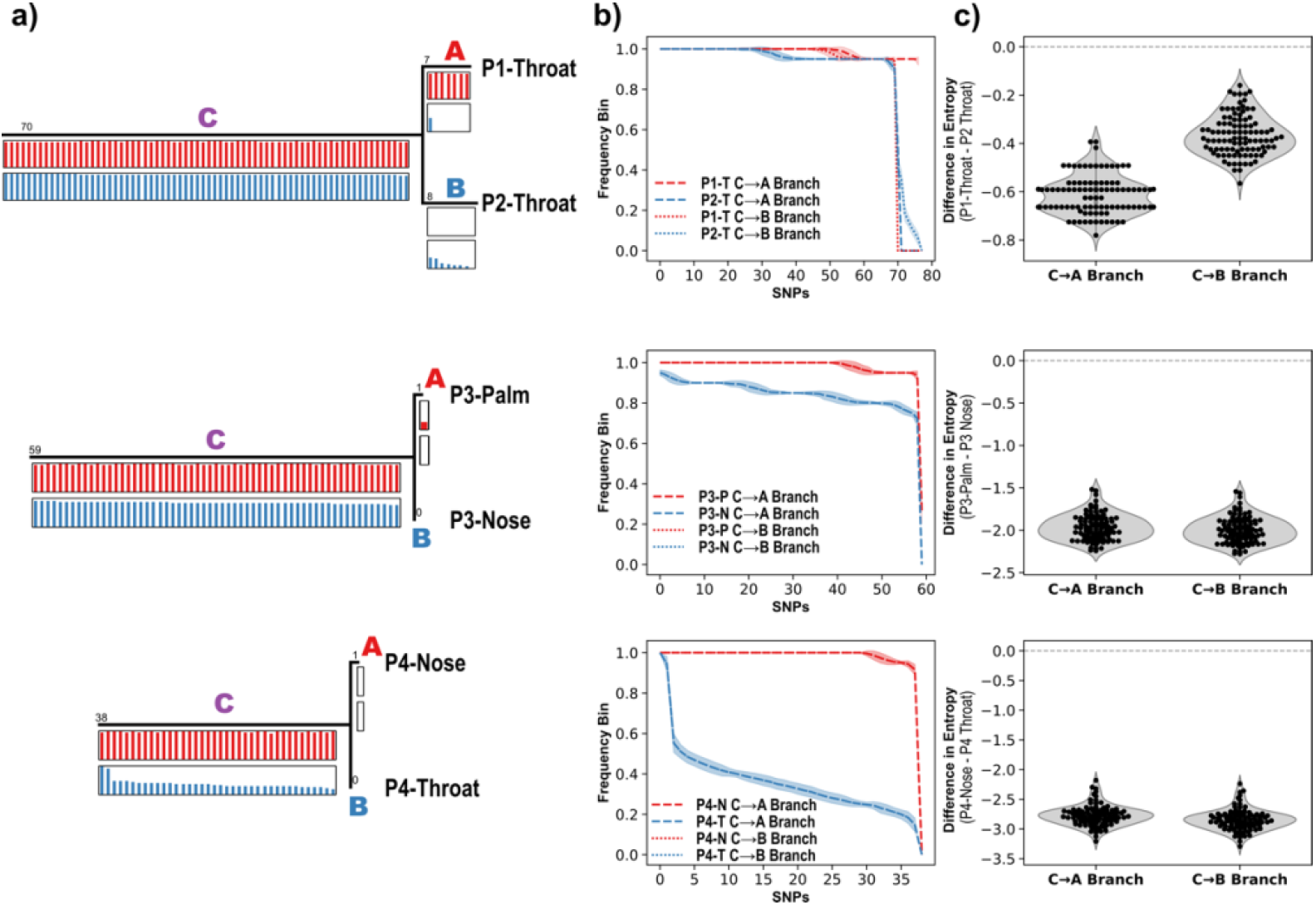
Population sequencing identifies a case of interhost transmission between two individuals and intrahost spread between different body sites. a) The trees of two clonal sequences and an outgroup (not shown) from throat samples from two individuals in a marital-type relationship living in the same home (top), a palm and nose sample from a single individual (middle), and a nose and throat sample from a single individual (bottom). Population sequencing from the samples provided the allele frequencies for the corresponding SNPs in the tree. b) To detect changes in allele frequencies along each branch, the frequencies were placed into bins (n=20) and Shannon entropy was calculated for the number of values in each bin. The SNP frequencies were resampled and re-binned (n=100) to estimate the entropy confidence interval. The plots show the binned frequency outcomes and standard deviation from resampling. c) The difference in resampled entropy estimates between each pair of populations along the C→A branch and the C→B branch. The negative values indicate that in each case, and on each branch, the second population had higher entropy than the first population which suggests that individual P2 was the transmission source for individual P1 (top), the nose was the source of spread to the palm in individual P3 (middle), and the throat was the source for the nasal population in individual P4 (bottom).

## Discussion

The high-throughput and low costs of second generation short-read sequencing technologies enable sequencing multiple individual samples to better describe the diversity of a population. Nonetheless, costs and sample preparation still limit the number of samples from a population that can be routinely sampled. The diversity within a single run will yield variants within the sequencing reads, however linking variants for haplotype reconstruction to describe the phylogenetic diversity of the sequenced population has only been successful in viral populations where a high density of variants enable reads to be stitched together [16]. Long-read sequencing technologies eliminate the need to stitch reads together in the smallest genomes, however these technologies are not likely to solve this problem of haplotype reconstruction in heterogeneous bacterial populations. Characterizing the diversity within a single population in a single sequencing run presents an important milestone in the development of tools that will revolutionize our understanding of microbial epidemiology. In this work, we have provided a theoretical framework through which a few sequencing runs can capture the phylogenetic diversity of an entire population. We tested this method with computer simulations to better understand limitations, and finally, apply this method to empirical examples to demonstrate the epidemiological utility.

### Population sequencing for characterizing genetic diversity

Population sequencing to characterize genetic diversity provides much potential for uncovering important epidemiological, ecological, and pathological insights associated with bacterial symbiotic relationships. We analyzed 20 isogenic clone sequences from 2 sputum samples and 2 population sequencing from one of the sputum samples collected on the same day during the 22nd year of chronic carriage of *B. pseudomallei* from a unique case of melioidosis. This analysis validates the utility of population sequencing in characterizing existing genetic diversity with a caveat that revealed a surprising aspect of this unique lung infection with implications on experimental design and potential insights into other pulmonary infections. The two population sequences contained variants that could be placed on different parts of the phylogeny. They also contained novel SNP mutations, indicating the existence of genetic diversity that is not observed among the 20 clones. However, the population samples did not capture all the phylogenetic diversity observed in the clones. In fact, the two population sequences themselves show almost no phylogenetic overlap, despite being derived from the same sputum sample. The structuring of the population sequences is consistent with the clonal samples and that the sputum was sampled as coughed up by the patient and not mixed or homogenized, suggesting bacterial population structure within the lungs themselves and the ability of sputum samples to maintain this structure. Therefore, the fact that the population sequences do not completely capture the phylogenetic diversity of the clones is due to the highly structured nature of the lung infection rather than a direct limitation of population sequencing. We had previously shown that in the 4th year of this infection there was no evidence of structuring [4], however our current results may be due to partitioning of lineages that developed in subsequent years or structuring at a much finer spatial scale than what was previously sampled. Other pathogens have been shown to exhibit spatial structuring in the lungs [10,35], and our results suggest that in some cases, non-invasive sputum sampling (and population sequencing) can be used as an alternative to bronchoscopy-based sampling to detect spatial structuring and, with attention to experimental design, to capture the genetic diversity. In fact, it is also possible that sputum-based sampling may be superior to other methods in detecting structuring at very small spatial scales. Our understanding about the diversity of different bacteria in various types of symbiotic relationships, in different parts of the body, and over time is in its infancy, but population sequencing certainly has the potential to accelerate research that unveils patterns of this important foundation of evolution.

### Population sequencing for determining directionality of transmission and spread

Determining the directionality of transmission and dissemination is often based on temporal epidemiological data. However, in many situations (e.g., epidemiological or forensic source attribution, for colonizing bacteria with a high prevalence in the population, species that cause chronic infections or have a delayed onset or absence of disease), the chronological order of disease onset or discovery of cases or forensic evidence may not be correlated with pathogen acquisition, making temporal based inferences unreliable and requiring phylogenetic analyses. However, phylogenetic analyses present challenges for characterizing person to person transmission and spread from one body site to another on a single person. In such cases, indicative phylogenetic signals may be limited by insufficient evolutionary time or confounded by too much evolutionary time and the transfer of a large portion of the population. As expected, our modeling showed a positive relationship between time before transmission and the likelihood of correctly determining directionality of transmission. This is because a lack of genetic diversity in the source population would result in two monophyletic sister clades after transmission causes a physical separation before additional evolution. As such, neither clade would be nested within the other and neither population would be expected to consistently harbor more genetic diversity. Conversely, increasing evolutionary time after transmission reduces the ability to determine transmission directionality. This is because an increase in diversity among the recipient population decreases the chances that greater diversity would be observed in the source population. Also as expected (Figure 3b-c), our modeling showed that a wider transmission bottleneck (when more cells are transferred) complicates our ability to determine directionality. This is also because the observed phylogenetic diversity in the recipient population will not be consistently less than the source population. These limitations can be ameliorated by selecting more clones for sequencing although selecting >5 clones is likely to present diminishing returns. While we know a little about infectious and lethal doses for a few pathogens, nothing is known about population sizes at the time of ecological establishment. Also, for most symbiotic bacteria, we know little about whether there are limitations on carriage time and population size before transmission or spread. Our simulations are therefore not designed to utilize known parameters, but rather to explore the general trends of expected limitations. It is however important to point out that we included no model of natural selection, and genetic drift was only incorporated once the carrying capacity was reached. This means that when a population was founded and until it reached the carrying capacity, any new mutation was guaranteed to increase in frequency. As drift, rather than selection, is the most powerful evolutionary force in small populations, these parameters may reflect reality regarding selection, but not for drift. However, both evolutionary forces reduce diversity, particularly in the recipient population, making the hallmarks of transmission directionality more prominent.

To demonstrate the potential of population sequencing and phylogenetic analysis to determine directionality of transmission and spread of bacteria, we used existing samples collected as part of our efforts to better understand carriage and transmission of *S. aureus*. Our example of transmission between people is interesting because it involves two individuals in a marital-like relationship. In such cases, close and frequent contact would be expected to result in a wide bottleneck and frequent transmission. Such conditions are expected to be particularly challenging for deciphering transmission directionality, and as such, we anticipated that entropy differences between the source and recipient populations would be minimal, however the entropy was greater for the population from the second person on each phylogenetic branch, providing a strong signal of transmission directionality. Our examples of spread between body sites of a single person (from the nose to the palm in Person 3 and from the throat to the nose in Person 4) involve similarly strong signals of directionality. Transmission from nose to palm is not surprising as the anterior nares is thought to be a principal reservoir for *S. aureus*. For this reason, our example showing spread from the throat to nose is particularly noteworthy and consistent with evidence that the throat may be an important reservoir. Longitudinal sampling and population sequencing from many people will be important for establishing the generalizability of these patterns that we observed across 4 individuals in characterizing carriage, sources, reservoirs, and transmission/spread of *S. aureus*.

## Methods

### Genotyping a population: Theoretical background

Genomic replication generates mutations that are inherited by subsequent generations. Single nucleotide polymorphisms (SNPs) are commonly used for phylogenetic inference because their low mutation rates make them relatively evolutionarily stable. As such, a SNP in one lineage is not likely to occur in another lineage over the course of a restricted evolutionary time frame [1]. Homoplasies in SNP loci (loci that revert to their original allelic state or mutate independently in different lineages) are rare but increase in quantity with evolutionary time. As a result, for short time-scale evolution (where bacteria follow a clonal model of inheritance), a single SNP can define a lineage as all descendants will contain the derived allele while all others will contain the ancestral allele. The stability of SNPs makes them rare in a genome, but this disadvantage is overcome by sampling and comparing entire genomes where the large genomic landscape increases the likelihood that informative characters can be found. SNPs in a population are discovered if a genome with that SNP is sequenced and compared to another genome. The phylogenetic location of discovered SNP mutations will be on the direct connecting evolutionary pathway between compared genomes (Pearson et al. 2004). Therefore, by determining the allelic states of samples without whole genome sequence data, it is possible to know where the sample diverged from the known evolutionary pathway, even though phylogenetic patterns since this bifurcation point will be unknown [1]. Using whole genome sequencing of a select few samples to guide SNP discovery for assay development and subsequent genotyping of a wider sample set has been frequently used for phylogenetic inference when high sequencing costs or paucity of genetic material prohibited whole genome sequencing of all samples. Now, we can apply the same underlying theory to characterize genomic diversity and phylogenetic patterns of the population contained in a single sample by leveraging individual reads in a single sequencing run.

### Population sequencing to characterize genomic diversity and phylogenetic patterns: Theoretical background

Deep sequencing directly from the population, provides reads from multiple genomes in the specimen, providing a representative sampling of the population diversity within the specimen. By mapping reads from such population sequencing to SNPs discovered through comparisons of isogenic clonally derived genome sequences (single colony DNA preparations), fine-scale population-level information can be superimposed upon a simplified but robust phylogeny. This is similar to how canonical SNP genotyping assays provide a rapid qPCR-based determination of clade membership, thus providing high-throughput phylogenetic information for samples that are not or cannot be sequenced (Figure 5). While deep sequencing allows for the discovery of all SNPs in the specimen population, linkage of SNPs across different reads to reconstruct haplotypes is not possible, limiting phylogenetic inference among variants outside the bifurcation point from the known phylogenetic backbone. As such, the same principles of phylogenetic discovery bias described previously for more diverse populations [1,17] apply: 1) Only SNPs along the direct connecting evolutionary path between sequenced clones will provide phylogenetic information, 2) phylogenetic collapse of lineages to their bifurcation point from the known phylogeny will occur, providing highly accurate, albeit incomplete phylogenetic information, 3) the clones sequenced for SNP discovery impact phylogenetic inference.

**Figure 5:**
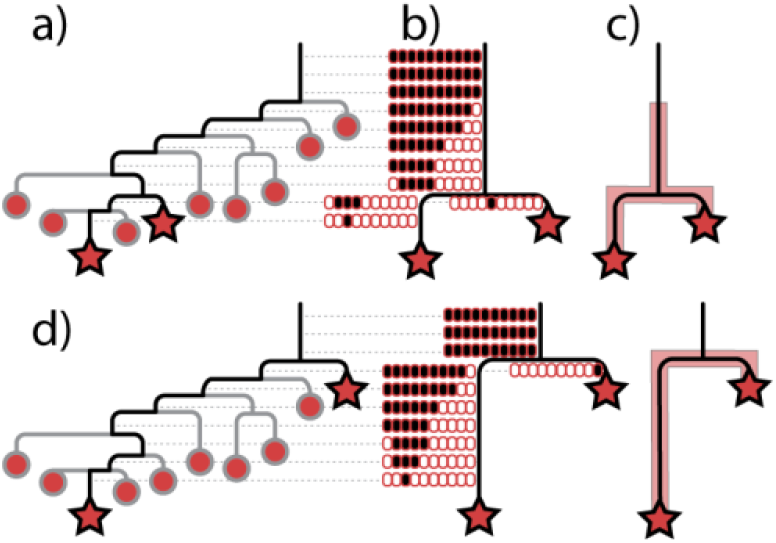
Inferring the phylogenetic distribution of a sequenced population with SNP discovery guided by two clonal sequences (stars) and an outgroup genome (not shown). a) Hypothetical phylogeny of all variants in a population. Stars represent sequenced clones for SNP discovery with SNPs being discovered along the direct connecting evolutionary branches (black branches). Determining the allele states for only the discovered SNPs will cause all other branches and variants (gray) to collapse upon the known phylogeny at their point of evolutionary divergence. b) A single sequence read (for simplicity) for the ten variants (ovals ordered from left to right according to the variants) in the population at each of the known SNP loci contain a black or white fill to represent the derived or ancestral (respectively) allele state. c) The allele states of the reads for SNPs along each known branch will be indicative of the phylogenetic boundaries of variants along these branches and can be represented as a semitransparent red cloud for this population. d) The phylogenetic position of sequenced clones will impact details of what is known about the phylogeny and distribution of the population, but not broader patterns.

### Empirical example characterizing genomic diversity and phylogenetic patterns: Genotyping a population of *Burkholderia pseudomallei* from a chronic lung infection

Patient 314 (P314) was diagnosed with melioidosis in the year 2000 and is the only known patient to have neither cleared nor succumbed to a *B. pseudomallei* infection over a more than two-decade period. We recently characterized and compared the phenotypes and genotypes of 118 isolates collected over the first >16 years of P314’s chronic carriage [4]. The evolutionary history of this population is well characterized with SNPs and small and large deletions accurately mapped onto the phylogeny. On August 30th, 2022, we collected two additional sputum samples from P314. From each of these samples, we directly plated sputum onto Ashdown’s agar (contains gentamicin and crystal violet) and also used a loop to cut away a small portion of the sputum that we stirred into Ashdown’s broth (contains colistin and crystal violet). After 2 days of broth incubation at 37°C, we plated the resulting biofilm on Ashdown’s agar. From the direct plating and plating following broth enrichment, we selected a composite of ten colonies (isogenic clones) from each sputum sample (20 total) for DNA extraction and genome sequencing. Also, from one of the sputum samples, we inoculated 25 mL and 4 mL of Ashdown’s broth) each with a piece of sputum to enrich for *B. pseudomallei* before DNA extraction (directly from broth and without plating) and sequencing. In summary, we sequenced 20 isogenic clones (from 2 sputum samples) and 2 populations (from 2 aliquots of one of the sputum samples). Sequencing was performed on the Illumina NextSeq using the P1 300 cycle kit. This work was approved by the Human Research Ethics Committee of the Northern Territory Department of Health and Menzies School of Health Research, Australia (ID HREC 02/38).

Genomic analyses, SNP discovery among clones and between a reference genome, and phylogenetic placement of clones was performed using previously described methods (Pearson et al. 2020). For genomic analysis of the two population sequences, we determined the percentage of reads for each allele of the 162 previously documented SNPs as well as 22 novel SNPs (leading to, and among the 20 new genomes). The population reads were preprocessed using a gatk best practices Snakemake (v7.19.1) [PMID: 34035898] workflow (https://github.com/snakemake-workflows/dna-seq-gatk-variant-calling/ v2.1.1) that performed quality control using fastqc v0.12.1 [18], read trimming using Trimmomatic v0.36 [19], read mapping using Samtools v1.12 [20] and BWA-MEM v0.7.17 [21] with default parameters against the MSHR1435 reference sequence (Accession numbers CP025264 and CP025265 [22]). LoFreq v2.1.5 [23] ‘viterbi’ command was used to perform probabilistic realignment of mapped reads and the ‘call’ function was used to call variants and estimate frequencies with a minimum coverage of 10 reads.

### Determining directionality of transmission/spread: Theoretical background

Transmission or spread of a pathogen occurs when a portion of the population in a source is transferred to another person, body site or environmental location. The introduced population faces attacks by the immune system or novel harsh environmental conditions, reducing the population size and genetic variation. While the infectious or lethal dose of some pathogens may be known, when this dose is greater than 1, we know of no examples where the initial size of the established population has been documented *in vivo*.

#### The simplest case

The initial established population may be as small as a single cell, representing a relatively simple transmission scenario where the diversity generated by descendants of this founding cell will all be contained within a monophyletic group nested within a paraphyletic source population (Figure 6a). Reads from sequencing can be used to determine the phylogenetic range of each population (Figure 6b), defined as positions along a branch where there is evidence that a derived allele is segregating (not fixed) in the population. The phylogenetic position of the clones selected for sequencing and SNP discovery frame what is known about branch lengths and the phylogenetic distribution of the populations (Figure 6b-d). In this most simple transmission scenario, the phylogenetic range of the population along the ancestral branch (henceforth referred to as Branch C), the branch leading to the sequenced clone from the source population (referred to as Branch A), and the branch to the sequenced clone from the recipient population (referred to as Branch B) will provide clues to the true nesting structure and the directionality of transmission. Here, we expect the following criteria to be true: 1) The recipient population will contain only derived alleles for SNPs along Branch C and only ancestral alleles for SNPs along Branch A. SNPs along Branch B may be segregating and thus contain a mixture of ancestral and derived alleles. These characteristics indicate that the phylogenetic range of the recipient population is limited to Branch B (Figure 6b-d). 2) In contrast, the source population may have segregating alleles at SNPs along Branch C (Figure 6b-c), indicating that the phylogenetic range of the population extends along Branch C. If the sequenced clone from the source population happens to be from the earliest diverging lineage, Branch C will contain fewer SNPs, all of which will be fixed for the derived allele (Figure 6d). 3) The source population will also have segregating alleles at SNPs along Branch A and possibly Branch B, depending on the phylogenetic position of the clone sequenced from the source population (Figure 6b-c). The presence of segregating alleles at SNPs along Branch C in one population and not the other is indicative of nesting and directionality of transmission (Figure 6b and c). Nesting and directionality can also be inferred if the source population has segregating alleles for SNPs along Branch B and the recipient population has no segregating alleles for SNPs along Branch A (Figure 6b and d).

**Figure 6:**
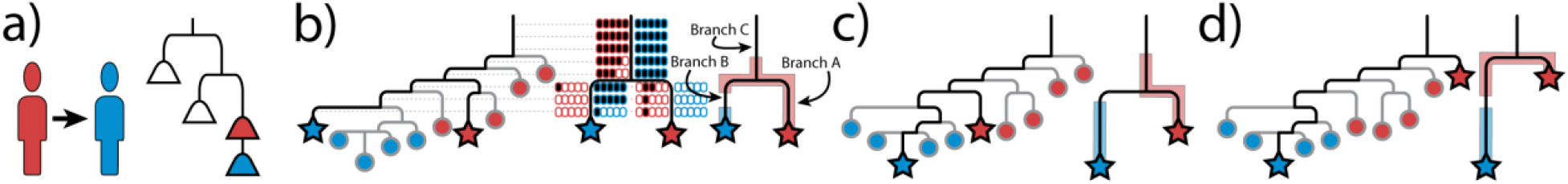
Determining directionality of transmission by sequencing a red and blue population guided by a clonal sequence from each population (stars) and an outgroup genome (not shown). a) Simple transmission scenario from red person to blue person involving an extreme bottleneck of a single cell to form the recipient (blue) population will result in the blue population forming a monophyletic clade nested within a paraphyletic source population. b) Hypothetical known phylogeny of all variants in the red and blue populations with stars represent sequenced clones for SNP discovery, ovals representing ancestral (white fill) and derived (black fill) alleles for each SNP along the corresponding branch and inferred phylogenetic range of each population (semitransparent colored areas) along the known phylogeny. c) and d) Consequences of different clones being sequenced and used for SNP discovery.

#### Closely related, but no evidence for transmission directionality

In cases where population sequencing does not provide evidence of transmission directionality, we expect no segregating alleles for SNPs on Branch C and neither population to have segregating alleles for SNPs on the terminal branch leading to the sequenced clone from the other population (Figure 7). This will be the case no matter which variants are sequenced for SNP discovery (Figure 7). The lack of phylogenetic evidence for transmission directionality does not exclude the possibility that transmission occurred as some variants may have gone extinct or not been sampled. Notably, proximity of the two population distributions provides evidence of epidemiological linkage that may include transmission despite the lack of evidence for directionality.

**Figure 7:**
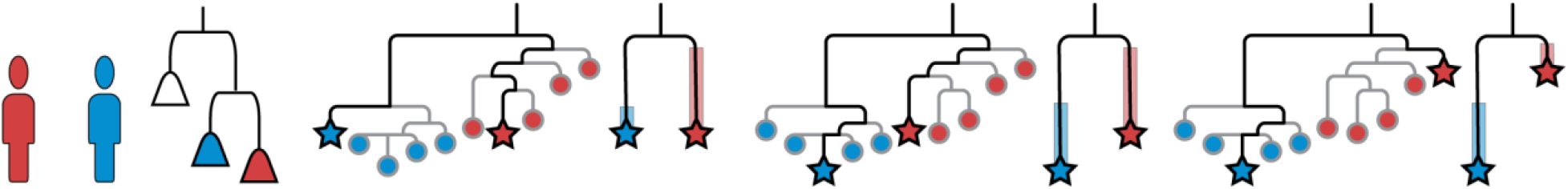
Population sequencing of two populations with no evidence of transmission. Both red and blue populations will be monophyletic and phylogenetic inference from population sequencing will show that the distribution and phylogenetic ranges of each population are restricted to the branch containing the sequenced clone used for SNP discovery. In this case, different sequenced clones have no impact on the distribution of populations being limited to Branches A or B.

#### More complex cases

More complicated transmission scenarios include multidirectional transmission (Figure 8a), multiple transmission events in the same direction, and transmission events with large bottlenecks where a larger portion of the source population becomes ecologically established in the recipient (Figure 8b-c). In these scenarios, the phylogenetic range of the source population may not always extend more ancestrally along Branch C, and the phylogenetic ranges of both populations along Branches A and/or B will often overlap. As greater diversity of the source is established within the recipient, overlap in phylogenetic range increases.

**Figure 8:**
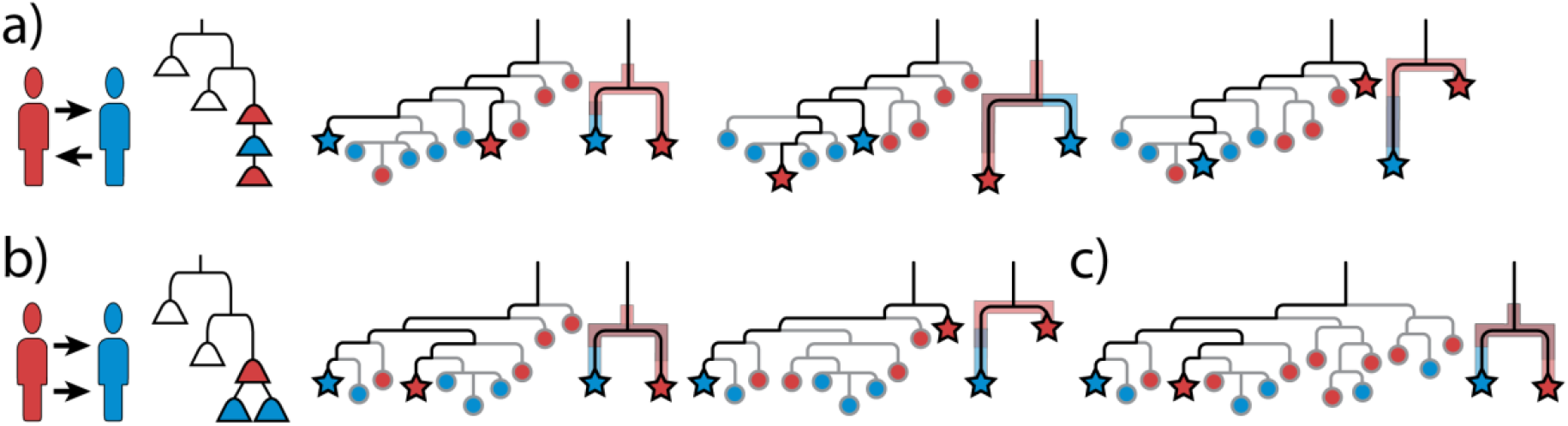
More complex transmission scenarios. a) Transmission from the red to blue person then back to the red person. Distribution of variants from population sequencing using various clones for SNP discovery show the red population being more ancestral. Unlike the simple transmission scenario shown in Figure 6, the phylogenetic range of variants in the red and blue population overlap with each other. b) Ongoing or multiple transmissions from red to blue resulting in the establishment of 2 clades in the recipient. The source population may have a more ancestral phylogenetic range along Branch C. As with the previous example, both populations have overlapping phylogenetic ranges. c) Transmission and establishment of 4 clades as might occur during multiple transmission events or a single event with a wide bottleneck. As more of the diversity is transferred from the source to the recipient, the populations can be expected to have increasingly overlapping ranges.

In complex transmission situations, consideration of only phylogenetic ranges will limit insights into population diversity and resulting inferences into phylogenetic nesting and transmission directionality (Figure 8c). However, sequence data also provides information on how portions of a population are distributed along lineages, allowing for comparisons of phylogenetic diversity to be used to determine phylogenetic nesting and transmission directionality. Even with a very wide bottleneck (when many cells are transferred and established in the recipient population), the recipient population will be less diverse than the source population with fewer lineages within each clade. This can be seen in the pattern in which allele frequencies change along a lineage in both populations (Figure 9). Each time a new lineage diverges, the proportion of the population (as measured in allele frequencies) along the lineage will decrease. The greater diversity in the source population means that there will be more diverging lineages and thus more changes in allele frequency than in the recipient population. The implication is that by comparing the changes in allele frequencies along the same lineage in each population, we can determine which population has greater diversity. To quantify the frequency changes along a lineage, we use a standard entropy metric. Similar frequencies are placed into bins and Shannon entropy [24] is calculated using the number of values in each bin normalized by the total number of frequencies using the following formula:

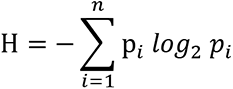

**Figure 9.**
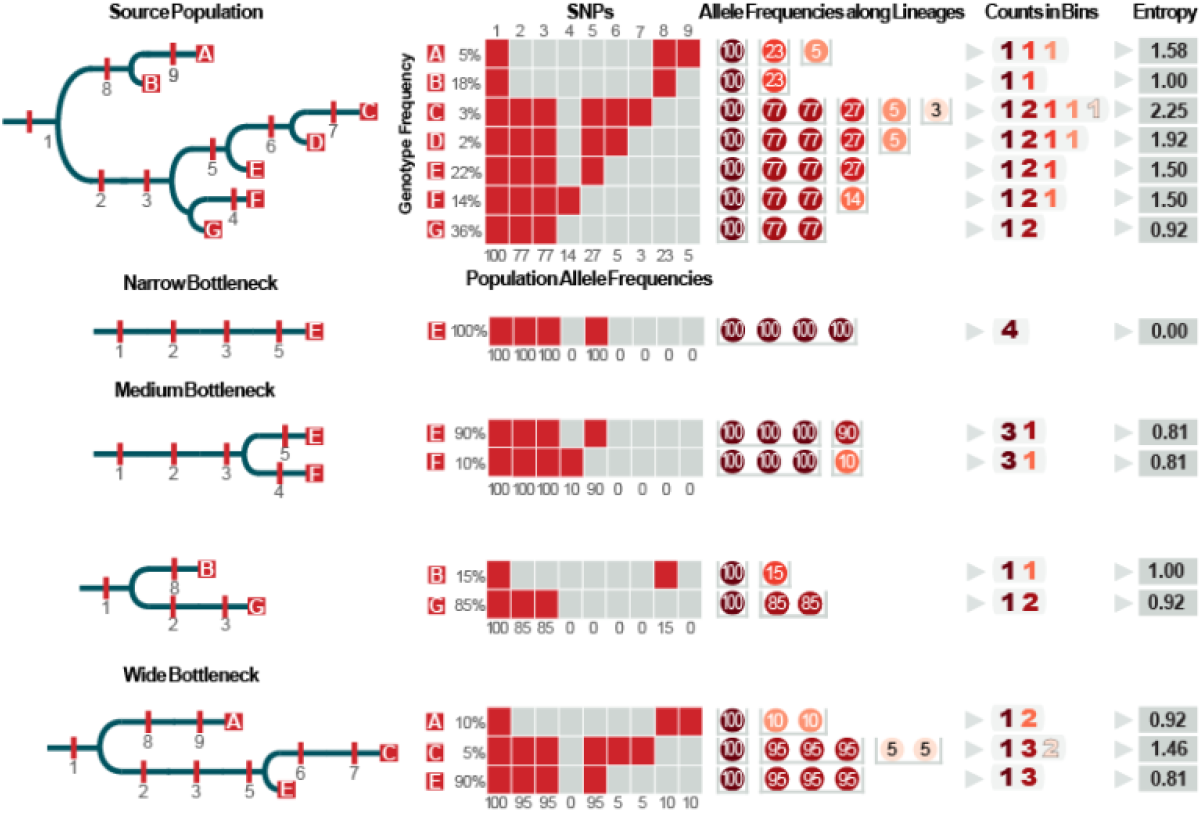
Comparing diversity using the entropy of allele frequencies along each lineage to determine transmission from a source to a recipient. In the diverse source population, the frequencies of the derived SNP alleles along a lineage will decrease each time a lineage diverges. When a transmission bottleneck occurs, there will be fewer diverging lineages which causes longer stretches of uninterrupted branches and runs of SNPs with same allele frequency. The pattern of allele frequency changes along a lineage can be measured and compared using a standard entropy formula. In a narrow bottleneck (only a single cell is transmitted), the allele frequencies of all SNPs will be the same (100%) and the entropy of the frequencies (0) is much lower than the same lineage in the source population (1.5). This pattern holds for wider bottlenecks, and while the entropy of allele frequencies for each lineage will usually be less than the entropy of the same lineage in the source population, it is theoretically possible for the entropy of the recipient population to be equal or even greater than the source if most of the population is transferred and concurrently eliminated from the source.

Where *n* is the number of bins, and *p_i_* is the number of frequencies in the *i^th^* bin divided by the total number of frequencies.

### Modeling to determine directionality of transmission/spread

We developed a computer simulation to model basic aspects of population evolution and transmission specifically to determine how population diversity is likely to impact our ability to discern transmission directionality. We did not employ precise empirical parameters from any specific organism or include more complex evolutionary processes. Our intent was not to understand how the evolutionary processes of mutation, natural selection, and drift, or different growth rates can impact diversity, but rather to discern how diversity (before, after, and during transmission) impacts our ability to determine directionality of transmission or spread. Because the impact of different evolutionary processes on the development of diversity is well-known, we should be able to extrapolate the impact of these processes on a case-by-case basis on our ability to determine directionality of transmission and spread. Detailed explanations of these simulations follow.

### Simulation

Populations were allowed to evolve from a single member into a heterogeneous group of members up to a carrying capacity of 100,000 over many generations. A population was modeled as a collection of genomes. A genome was modeled as a collection of mutations. A mutation was modeled as an integer in the range [0, 2,800,000]. The upper limit of 2,800,000 reflects the approximate size of a *S. aureus* genome, chosen because of our empirical examples of spread and transmission. A generation was modeled as a series of the following events: mutation, duplication, and pruning. Mutations were modeled each generation according to a Poisson distribution with an expected rate λ of 1.6 × 10^−10^ mutations per genome position per generation, which were then applied randomly to members of the population. Duplication modeled binary fission by duplicating each genome in the population. When duplication resulted in a population size greater than the carrying capacity, pruning randomly removed genomes from the population until a population size equal to the carrying capacity was restored. Pruning therefore modeled genetic drift that was only in effect at the carrying capacity. Selection was not incorporated into the simulation and all mutations were thus equally likely to persist. A source population was allowed to evolve first, for some specified number of generations. Then, transmission was modeled by randomly selecting a subset of the source population as founders to a recipient population. This process was parameterized by the modeled transmission bottleneck: the number of members (genomes) copied from the source population to the recipient population. The recipient population and the source population were then allowed to evolve independently for a specified number of generations.

### Analyses and calculating transmission directionality

After a simulation completed, a random sample of genomes was taken from both the source and recipient populations. The size of the sample was configurable but always equal for both populations. In cases where the population sizes were sufficiently large, sampling was performed without replacement. In cases where a small number of post-transmission generations resulted in a small recipient population, sampling with replacement was performed to meet our desired sample size.

Next, all possible pairs of genomes between the source and recipient populations were compared. A phylogenetic tree was constructed for each pair using the single nucleotide polymorphisms (SNPs) between the two genomes. This resulted in n = (sample size)^2^ trees. For each SNP locus, we determined the fraction of genomes in each population that contained each allele. In each population, SNP alleles were therefore either fixed (when all genomes in a population contain the same SNP allele) or segregating (when both alleles are present in a population), with the proportion of each allele in each population documented. Using this information coupled with the phylogenetic location of each SNP, we evaluated three methods of determining transmission directionality:

#### Method 1

phylogenetic range on the ancestral branch (Branch C): Because we assume greater diversity in the source population, the source population will often extend more ancestrally along Branch C. We therefore expect that the number of segregating SNPs on Branch C will be greater in the source population. We recorded the proportion of simulations where: 1) the source population had more segregating SNPs on Branch C than the recipient population — indicating correct determination of transmission directionality, 2) the recipient population had more segregating SNPs on Branch C than the source population — indicating incorrect determination of transmission directionality, and 3) the numbers of segregating SNPs on Branch C in each population were equal — indicating ambiguity in the directionality of transmission.

#### Method 2

phylogenetic range on Branches A and B: A more diverse source population will also likely extend along both Branches A and B. Conversely, a recipient with less diversity will have a restricted range along these two branches. We therefore recorded the proportion of simulations where: 1) the source population had more segregating SNPs along Branch B than the recipient population had on Branch A — indicating correct determination of transmission directionality, 2) the source population had fewer segregating SNPs along Branch B compared to the number of segregating SNPs in the recipient population along Branch A — indicating incorrect determination of transmission directionality, and 3) the source and recipient populations had an equal number of segregating SNPs along Branch B and Branch A, respectively — indicating ambiguity in the directionality of transmission.

#### Method 3

phylogenetic entropy: While the first two methods account for diversity by simply defining and comparing the phylogenetic ranges, they do not capture and leverage the distribution of each population along the branches. Assuming a more diverse source population, we expect the source population to have more bifurcation points (Figure 9). To estimate the likelihood of correctly determining transmission directionality, we recorded the proportion of simulations where the source had: 1) greater entropy of allelic frequency on both Branches C through A and Branches C through B, or 2) greater entropy on one of these lineages and equal on the other. Incorrect determination of transmission occurred when the recipient population had greater entropy in these situations, and ambiguous results occurred when entropy values were greater in the recipient population on one branch and greater in the source population on the other branch or were equal for each population along both branches.

### Empirical example of population sequencing of *S. aureus* to determine transmission directionality

We used samples from an ongoing research project designed to better understand community carriage and transmission among social groups in a population on the United States/Mexico border [25–29]. These samples were part of Project 1116783 that was approved by the Northern Arizona University Institutional Review Board. We used double-tipped BBLCultureSwab swabs to swab the nose, throat, and palms of participants. Swabs were stored on ice for no longer than 24 hours to maximize the likelihood of cell survival [30]. Each swab was streaked onto CHROMagar *S. aureus* media and incubated for 24 hours at 37°C. Colonies in the pink to mauve color range were considered to be *S. aureus*. One colony from each swab was streaked for isolation and sequenced to provide an isogenic clonal representative. All other suspected *S. aureus* colonies were collected, combined, and sequenced to represent the entire population. DNA extraction, library construction, and sequencing were performed as previously reported [28].

The clonal reads underwent quality control using fastp v0.20.1 [31] using default parameters and the SNPs were called using NASP v1.2.0 [32] which mapped the reads to the reference (accession NC_007795) using BWA-MEM v0.7.17 [21] and called the SNPs using the GATK v3.8 UnifiedGenotyper [33,34] method. SNP calls required a minimum coverage of 10 reads and at least 90% of the reads supporting the SNP call. The processing steps for the combined population samples were the same as described above for the *B. pseudomallei* populations.

## Acknowledgements

This work was funded by NAU Southwest Health Equities Research Collaborative (NIH/NIMHD U54MD012388) and two grants from NIH/NIAID (R01AI172924 and R15AI156771).

## Data Availability

The code for running and analyzing the simulations is archived at https://doi.org/10.5281/zenodo.11438885. The *B. pseudomallei* sequencing reads are available through NCBI’s Sequence Read Archive under Bioproject PRJNA321854. The *S. aureus* sequencing reads are available under Bioprojects PRJNA1111899 and PRJNA660486. The results from analyzing the *B. pseudomallei* and *S. aureus* samples are available at https://doi.org/10.6084/m9.figshare.25955911.v1.

